# Characterization of entry pathways, species-specific ACE2 residues determining entry, and antibody neutralization evasion of Omicron BA.1, BA.1.1, BA.2, and BA.3 variants

**DOI:** 10.1101/2022.06.01.494385

**Authors:** Sabari Nath Neerukonda, Richard Wang, Russell Vassell, Haseebullah Baha, Sabrina Lusvarghi, Shufeng Liu, Tony Wang, Carol D. Weiss, Wei Wang

## Abstract

The SARS-CoV-2 Omicron variants were first detected in November 2021, and several Omicron lineages (BA.1, BA.2, BA.3, BA.4, and BA.5) have since rapidly emerged. Studies characterizing the mechanisms of Omicron variant infection and sensitivity to neutralizing antibodies induced upon vaccination are ongoing by several groups. In the present study, we used pseudoviruses to show that the transmembrane serine protease 2 (TMPRSS2) enhances infection of BA.1, BA.1.1, BA.2, and BA.3 Omicron variants to lesser extent compared to ancestral D614G. We further show that Omicron variants have higher sensitivity to inhibition by soluble angiotensin converting enzyme 2 (ACE2) and the endosomal inhibitor chloroquine compared to D614G. The Omicron variants also more efficiently used ACE2 receptors from nine out of ten animal species tested, and unlike the D614G variant, used mouse ACE2 due to the Q493R and Q498R spike substitutions. Finally, neutralization of the Omicron variants by antibodies induced by three doses of Pfizer/BNT162b2 mRNA vaccine was 7-8-fold less potent than the D614G, and the Omicron variants still evade neutralization more efficiently.

## Introduction

Since the origin of the COVID-19 pandemic, several SARS-CoV-2 variants of concern (VOCs) with enhanced transmissibility, ACE2 binding affinity and immune evasive properties have emerged, including the most recent Omicron VOCs. Currently, Omicron (B.1.1.529) is comprised of five main lineages designated BA.1 (and its sublineage BA.1.1), BA.2 (and its sublineage BA.2.12.1), BA.3, BA.4, and BA.5. BA.1 was first identified in November 2021 in Botswana, and it rapidly replaced the then dominant Delta (B.1.617.2) VOC to become globally prevalent due to its enhanced transmissibility and ability to evade antibody neutralization ^1–6^. By early 2022, an alarming rise of BA.2 was seen in several parts of the world, leading to the replacement of BA.1 and BA.1.1. In comparison to BA.1, BA.2 was demonstrated to have faster replication kinetics, enhanced fusogenicity, and greater pathogenicity in hamsters ^7,8^. BA.4 and BA.5 are two new lineages that are presently emerging in South Africa.

The spike protein of Omicron variants bears an unprecedented degree of antigenic divergence with the highest number of substitutions compared to ancestral B.1 (Wuhan-Hu-1, and D614G) variants and earlier VOCs. These include 21 spike substitutions shared by the three main lineages of Omicron in the N-terminal domain (NTD) (G142D), receptor binding domain (RBD) (G339D, S373P, S375F, K417N, N440K, S477N, T478K, E484A, Q493R, Q498R, N501Y, Y505H), and furin cleavage site proximity (D614G, H655Y, N679K, P681H) of the S1 subunit, as well as substitutions in fusion peptide proximity (N764K, D796Y) and heptad repeat region 1 (HR1) (Q954H, N969K) of the S2 subunit (Figure 1). Additionally, 16 unique insertions/deletions/substitutions in BA.1 and 9 unique insertions/substitutions in BA.2 are present. BA.3 shares 10 unique substitutions/deletions with BA.1 and two unique substitutions with BA.2. BA.1.1 differs from BA.1 by one RBD substitution (R346K).

**Figure 1.**
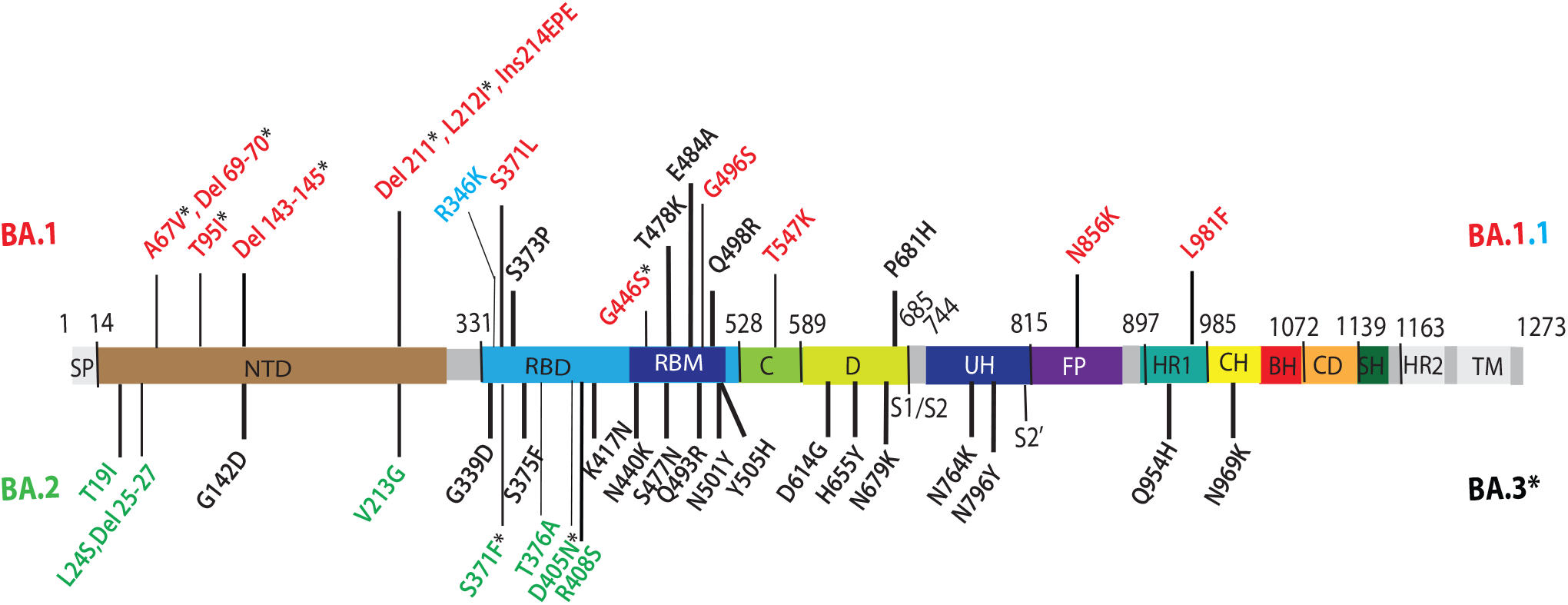
Omicron lineage substitutions in spike. Omicron substitutions are shown in a primary structure of the SARS-CoV-2 spike protein, with various domains and cleavage sites indicated. SP, signal peptide; NTD, N-terminal domain; RBD, receptor binding domain; RBM, receptor binding motif; C, C domain; D, domain D; S1/S2, furin cleavage junction of S1/S2 subunits; UH, upstream helix; FP, fusion peptide; HR1/2, heptad repeat 1/2; CH, central helix; BH, beta hairpin; CD, connector domain; SH, stem helix; TM, transmembrane domain. Substitutions common to Omicron (BA) VOCs are shown in black. Substitutions unique to BA.1 VOC are shown in red. R346K substitution additionally present in BA.1.1 is shown in light blue. Substitutions unique to BA.2 VOC are shown in green. Substitutions in BA.1 and BA.2 shared by BA.3 VOC are shown as residues marked with an asterisk.

Several reports continue to demonstrate total loss of or severely dampened neutralizing activity of serum or plasma obtained from previously convalesced individuals, recipients of two doses of the Pfizer–BioNTech BNT162b2 or Moderna mRNA-1273 vaccine, as well as several therapeutic monoclonal antibodies against the BA.1 and BA.2 VOCs ^1,3,9–14^. A third booster dose of either Pfizer–BioNTech BNT162b2 or Moderna mRNA-1273 vaccine, however, recalls and expands preexisting antigen specific memory B cell clones and generates novel B cell clones resulting in enhanced neutralizing antibody titers and breadth towards BA. 1 and BA.2 VOCs ^15,16^. Booster-elicited neutralizing antibody titers remain durable for at least 4 months ^17,18^.

Apart from humans, SARS-CoV-2 was also found to naturally infect diverse domestic and wild animal species, including farm minks ^19,20^, companion pets (e.g., cats, dogs, ferrets, Syrian hamsters) ^21–23^, zoo animals (e.g., lions, tigers, cougars, snow leopards, gorillas, otters, and hippopotami) ^24^, and free ranging white-tailed deer ^25^. Furthermore, experimental infections in livestock species have determined low-level replication of ancestral variants in cattle ^26,27^, pigs ^28^, and sheep ^27,29^. These observations signify the broad host range of SARS-CoV-2 and the risk of infection by heavily mutated variants to give rise to potential reservoirs that may pose a further risk for spillover back to humans.

The primary genetic determinant of SARS-CoV-2 host range is spike interaction with species-specific receptor to allow subsequent viral entry. SARS-CoV-2 enters host cells by interacting with angiotensin-converting enzyme 2 (ACE2) in a species-specific manner. For instance, ancestral SARS-CoV-2 spike does not interact with murine ACE2, and therefore, human ACE2 (hACE2) transgenic murine models ^30–32^ or mouse-adapted SARS-CoV-2 ^33^ were developed to study SARS-CoV-2 pathogenesis. The trimeric spike is a class I fusion protein that is cleaved into the S1 and S2 subunits, which are noncovalently associated on the surface of virions. Following the interaction of the RBD (residues 331 to 528) of the S1 subunit with the N-terminus of ACE2 and further proteolytic processing of the S2 subunit at the S2’ site by the host proteases (cathepsin L in endosomes, or TMPRSS2 on the plasma membrane), extensive and irreversible conformational changes occur in the S2 subunit to facilitate membrane fusion. The insertion of S2 fusion peptide in the target cell membrane and the interaction between HR1 and HR2 of the S2 subunit result in the formation of a stable six-helix bundle that brings the viral and cell membranes into proximity for fusion and subsequent viral entry.

Here, we report that compared to ancestral D614G, Omicron variants’ (BA.1, BA.2, BA.3, and BA.1.1) infection is less influenced by TMPRSS2, and Omicron variants are therefore relatively more susceptible to endosomal entry inhibition. Additionally, Omicron variants exhibit greater sensitivity to soluble ACE2 (sACE2) inhibition. Furthermore, Omicron variants’ spikes have more efficient usage of ACE2 orthologs from nine diverse animal species than D614G. In addition, we found that Q493R and Q498R substitutions in the Omicron variant spikes promote usage of mouse ACE2 for entry whereas the Q493R substitution prevents usage of Chinese rufous horseshoe bat ACE2. Finally, sera obtained from fully vaccinated (three doses of the Pfizer/BNT162b2 vaccine) individuals and fully vaccinated individuals with a breakthrough infection of Omicron potently neutralize pseudoviruses bearing spike proteins of D614G and Omicron variants. However, neutralization titers against the Omicron variants are 7-8-fold lower compared to D614G.

## Results

### Infectivity and endosomal entry of Omicron variant pseudoviruses

Pseudoviruses bearing spike proteins of BA.1, BA.2, BA.3, and BA.1.1 Omicron variants displayed similar infectivity as D614G in 293T.ACE2 and 293T.ACE2.TMPRSS2 cells (Figure 2A). The presence of TMPRSS2 resulted in a 2.4-14.2-fold enhancement of infection for all the pseudoviruses in 293T.ACE2.TMPRSS2 cells compared to 293T.ACE2 cells. However, D614G pseudovirus infection was greatly enhanced (14.2-fold;*p≤0.0001*) as observed previously ^34^, whereas infection of Omicron variants was comparatively less enhanced (2.4-4.9-fold; *p≤0.0001*) in the presence of TMPRSS2 (Figure 2A and 2B). These findings indicate that TMPRSS2 confers relatively less enhancement of Omicron entry into cells than D614G.

**Figure 2.**
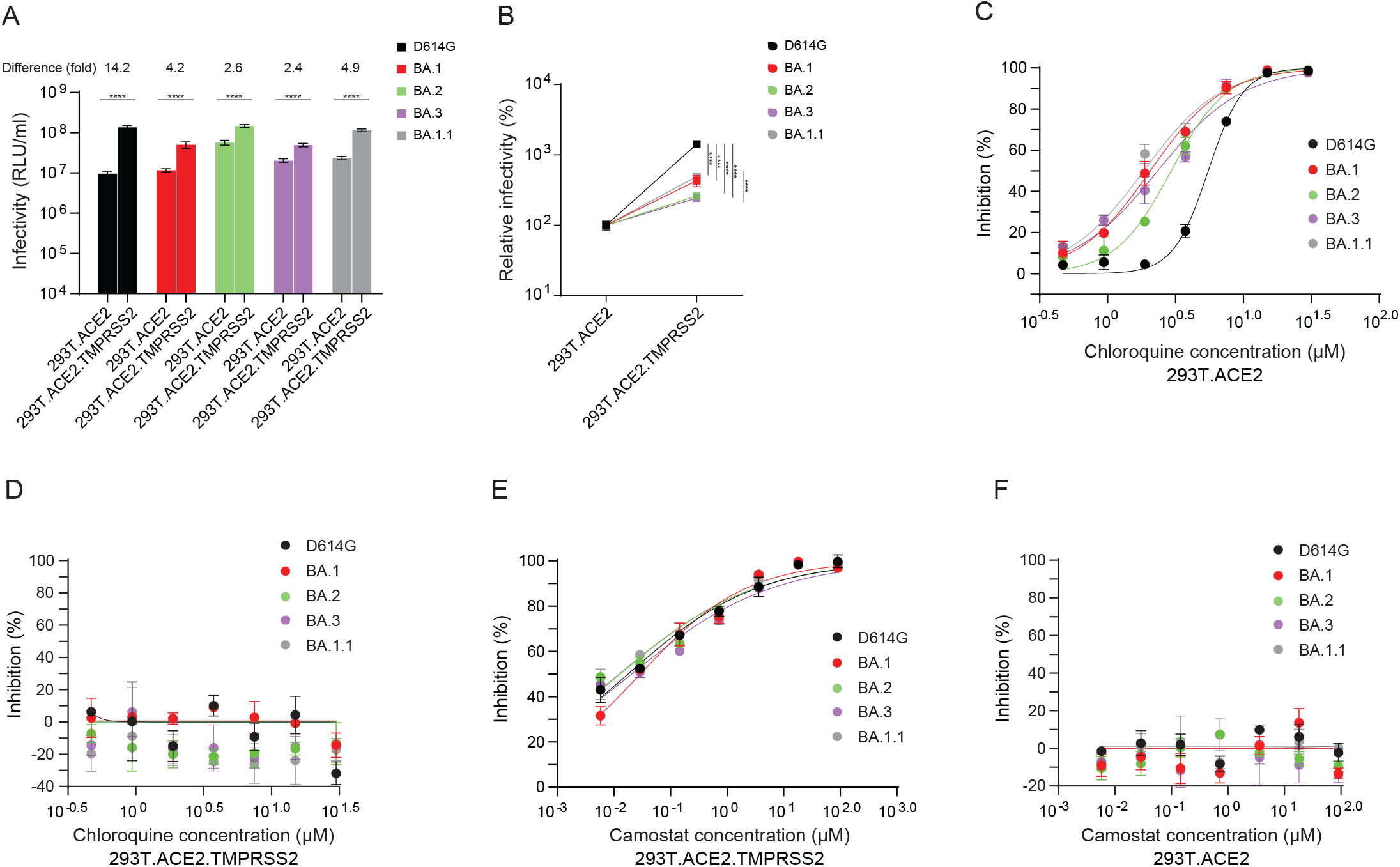
Infection and cell entry mechanisms of Omicron lineage pseudoviruses. Infectivity of D614G and Omicron (BA) lineage pseudoviruses on 293T.ACE2 and 293T.ACE2.TMPRSS2 cells. Each cell type was infected with D614G and Omicron lineage pseudoviruses **(A)**. Relative infectivity of D614G and Omicron lineage pseudoviruses in stable 293T.ACE2 cells compared to stable 293T.ACE2.TMPRSS2 cells **(B)**. Chloroquine sensitivity of D614G and Omicron lineage pseudoviruses in stable 293T.ACE2 **(C)** and 293T.ACE2.TMPRSS2 cells **(D)**. Cells were pretreated with indicted concentration of chloroquine for 2 h prior to infections with pseudoviruses in media containing the inhibitor. Camostat sensitivity of D614G and Omicron lineage pseudoviruses in stable 293T.ACE2.TMPRSS2 **(E)** and 293T.ACE2 cells **(F)**. Cells were pretreated with indicted concentration of chloroquine for 2 h prior to infection with pseudoviruses in media containing the inhibitor. X-axis indicates concentration of inhibitor. Y axis indicates percentage inhibition compared to pseudovirus infection without inhibitor treatment. Results shown are representative of three independent experiments. Asterisk* denotes significance: *:*p ≤ 0.05;* **: *p ≤ 0.01;* ***:*p ≤ 0.0005;* ****:*p ≤ 0.0001*.

Several recent studies showed delayed or attenuated replication of BA.1, BA.2, and BA.1.1 variants compared to B.1 and Delta VOCs in TMPRSS2-expressing cell lines (Calu-3, Caco-2, VeroE6/TMPRSS2) compared to VeroE6 cells, along with higher sensitivity to endosomal inhibitors (Chloroquine, Bafilomycin A) and less sensitivity to TMPRSS2 inhibitor (Camostat mesylate) ^6,8,35,36^. In cells expressing ACE2 but not TMPRSS2, spike-mediated entry follows the endosomal route, whereas in cells expressing both ACE2 and TMPRSS2, spike-mediated entry occurs mainly at the cell surface. We investigated ACE2 preference and TMPRSS2 usage by examining the sensitivity of D614G and the Omicron (BA.1, BA.2, BA.3 and BA.1.1) variant pseudoviruses to endosomal and TMPRSS2 inhibitors, chloroquine and camostat mesylate, respectively. In 293T.ACE2 cells, chloroquine potently inhibited viral entry of both D614G and Omicron variants, but Omicron pseudoviruses were more sensitive to chloroquine (BA.1 IC_50_:2.06,*p<0.001;* BA.2 IC_50_:2.94,*p<0.005;* BA.3: 2.34,*p<0.001;* BA.1.1 IC_50_: 1.85,*p<0.002*) than D614G (IC_50_: 5.51) (Figure 2C). No chloroquine inhibition was observed in 293T.ACE2.TMPRSS2 cells (Figure 2D), suggesting that the presence of TMPRSS2 facilitates entry via the cell surface for all variants. D614G (IC_50_: 0.02) and Omicron (BA.1 IC_50_: 0.03; BA.2 IC_50_: 0.01; BA.3: 0.02; BA.1.1 IC_50_: 0.01) pseudoviruses displayed similar sensitivity to camostat mesylate in 293T.ACE2.TMPRSS2 cells (Figure 2E). These findings indicate greater sensitivity of Omicron pseudoviruses to endosomal entry inhibition compared to D614G.

### Sensitivity of Omicron variants to soluble ACE2 neutralization

We further investigated ACE2 binding of Omicron variants by analyzing the sensitivity of D614G and Omicron pseudoviruses to soluble human ACE2 (sACE2) in a neutralization assay. Compared to D614G (IC_50_: 3.31 μg/ml), the BA.1 (IC_50_: 0.98μg/ml,*p<0.0001*), BA.2 (IC_50_: 0.99 μg/ml, *p<0.0004*) and BA.1.1 (IC_50_: 0.99 μg/ml, *p<0.0005*) variants demonstrated approximately 3-fold greater sensitivity to sACE2 whereas the BA.3 variant (IC_50_: 1.85 μg/ml, *p<0.0005*) showed 1.75-fold greater sensitivity to sACE2 (Figure 3). The sACE2 IC_50_ values reported here are comparable to previously reported values for D614G and BA.1 ^37^.

**Figure 3.**
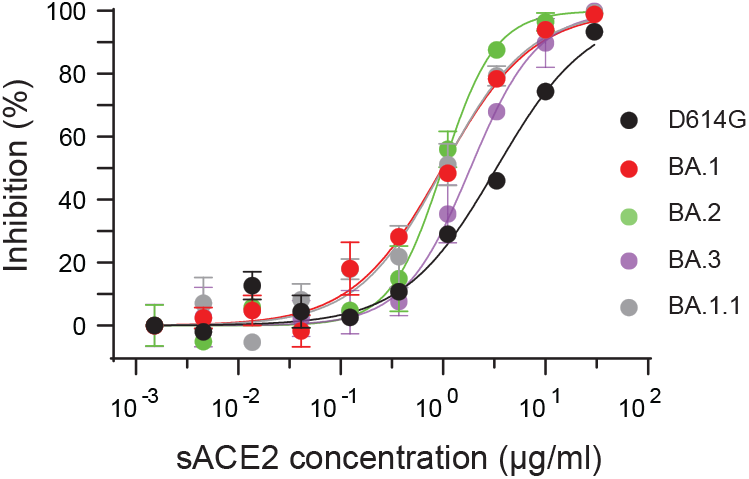
The neutralization of soluble ACE2 to Omicron lineage pseudoviruses. Sensitivity of D614G and Omicron lineage pseudoviruses to soluble human ACE2 was evaluated on stable 293T.ACE2.TMPRSS2 cells. Results shown are representative of three independent experiments.

### Pseudovirus stability of Omicron variants at various temperatures

Pre-incubation of viruses at different temperatures can reveal differences in the stability of the viral glycoprotein trimers, including SARS-CoV-2 ^37,38^. We evaluated the effect of temperature on spike glycoprotein stability, and thus infectivity, by incubating D614G and Omicron variants pseudoviruses at 4°C, 25°C (room temperature or RT), 32°C, 37°C, 42°C and 50°C for an hour or 50°C for various periods of time before measuring their infectivity on 293T.ACE2 and 293T.ACE2.TMPRSS2 cells. We observed no significant differences in the infectivity of D614G and Omicron variants pseudoviruses after incubation at 4°C, RT, 32°C, 37°C, and 42°C (Figure 4). All viruses were relatively stable at 4°C, RT, 32°C, 37°C, and 42°C. Our findings are in agreement with a previous study where no differences in infectivity were observed between VSV-pseudotyped D614G and BA.1 VOC pseudoviruses upon extended incubation (72hrs) at 4°C, RT, and 37°C ^37^. However, infectivity of all pseudoviruses dramatically declined upon incubation at 50°C with no significant difference between D614G and Omicron variants pseudoviruses (Figure 4).

**Figure 4.**
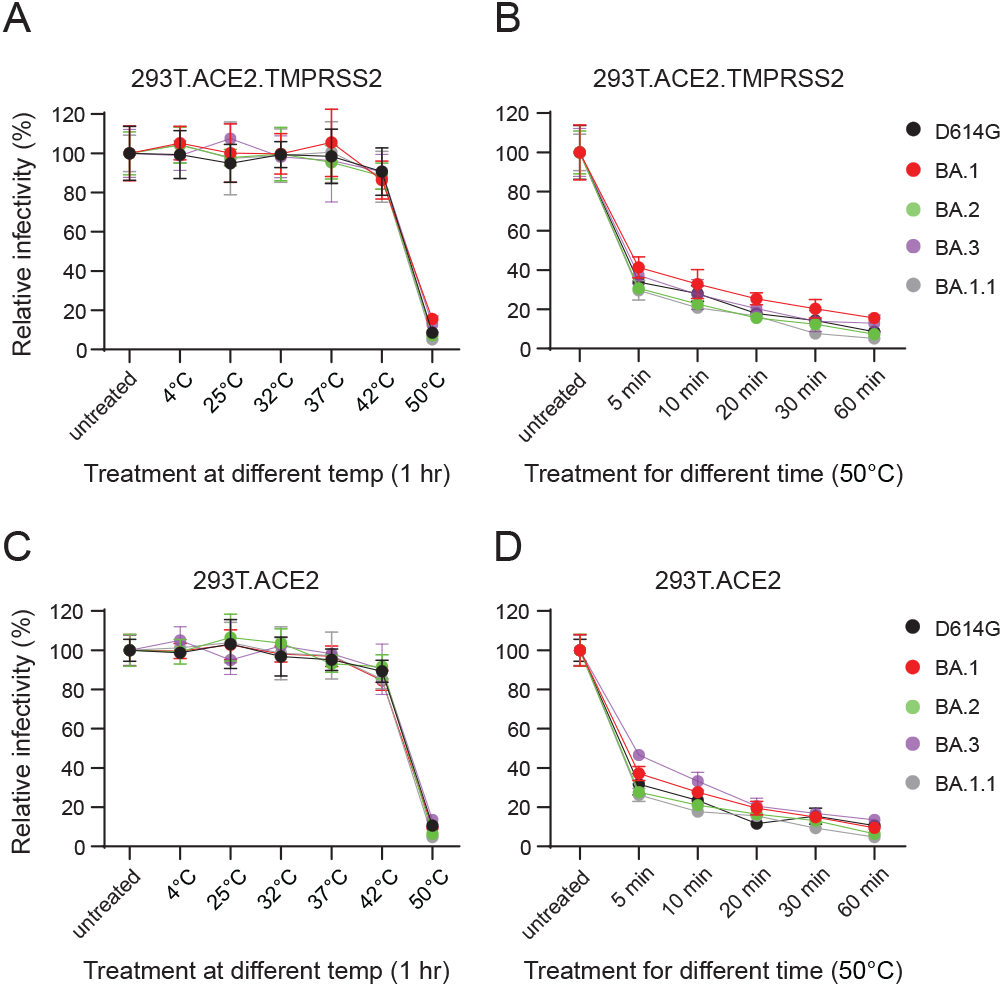
Omicron lineage pseudoviruses stability. Temperature stability of D614G and Omicron (BA) lineage pseudoviruses on 293T.ACE2.TMPRSS2 and 293T.ACE2 cells were evaluated. Pseudoviruses were untreated or subjected to various temperatures for an hour prior to infectivity on 293T.ACE2.TMPRSS2 (**A**) and 293T.ACE2 (**C**) cells. Pseudoviruses were untreated or subjected to 50°C for different durations prior to infectivity on 293T.ACE2.TMPRSS2 (**B**) and 293T.ACE2 (**D**) cells. Results shown are representative of two independent experiments.

### Omicron variants display distinct ACE2 receptor usage compared to D614G

Substitutions in the RBD have been linked to SARS-CoV-2 adaptation to new hosts, such as ferrets ^39^, mouse ^33^, mink ^40^ and white-tailed deer ^41^. The ongoing dominance and circulation of Omicron VOCs pose a significant risk for reverse zoonosis and spill back into the humans. Therefore, we investigated the ability of the Omicron (BA.1, BA.2, BA.3 and BA.1.1) variants to use ACE2 orthologs from ten diverse host species including African green monkey, Chinese rufous horseshoe bat, ferret, mouse, Chinese hamster, Syrian golden hamster, white-tailed deer, swine, bovine, and Malayan pangolin. 293T cells transiently transfected with ACE2 receptors of each species were infected with pseudoviruses bearing spike proteins of D614G and the Omicron variants. We found that D614G pseudoviruses infected cells expressing ACE2 receptors of all the species well above the background except mouse, consistent with prior studies ^3,33,42^ (Figure 5A). Conversely, ACE2 receptors of all species, except horseshoe bat, supported infection by Omicron pseudoviruses (Figure 5B-E). 293T cells with stable expression of human ACE2 (293T.ACE2) were used as a positive control. We confirmed robust ACE2 expression for each species via Western blotting using the V5 tag at the C-terminus of these ACE2 proteins (Supplementary Figure 1). The Omicron pseudoviruses had significantly higher levels of infection than D614G pseudoviruses in cells expressing African green monkey, Chinese rufous horseshoe bat, ferret, mouse, Chinese hamster, Syrian golden hamster, white-tailed deer (save BA.1), swine, and bovine ACE2 receptors (Table 1). No significant difference between D614G and Omicron variants was observed with respect to Malayan pangolin ACE2 (Table 1). These findings indicate that the receptor binding motif (RBM) substitutions in the Omicron (BA.1, BA.2, BA.3 and BA.1.1) variants can modulate ACE2 receptor usage and therefore, Omicron variants’ entry.

**Figure 5.**
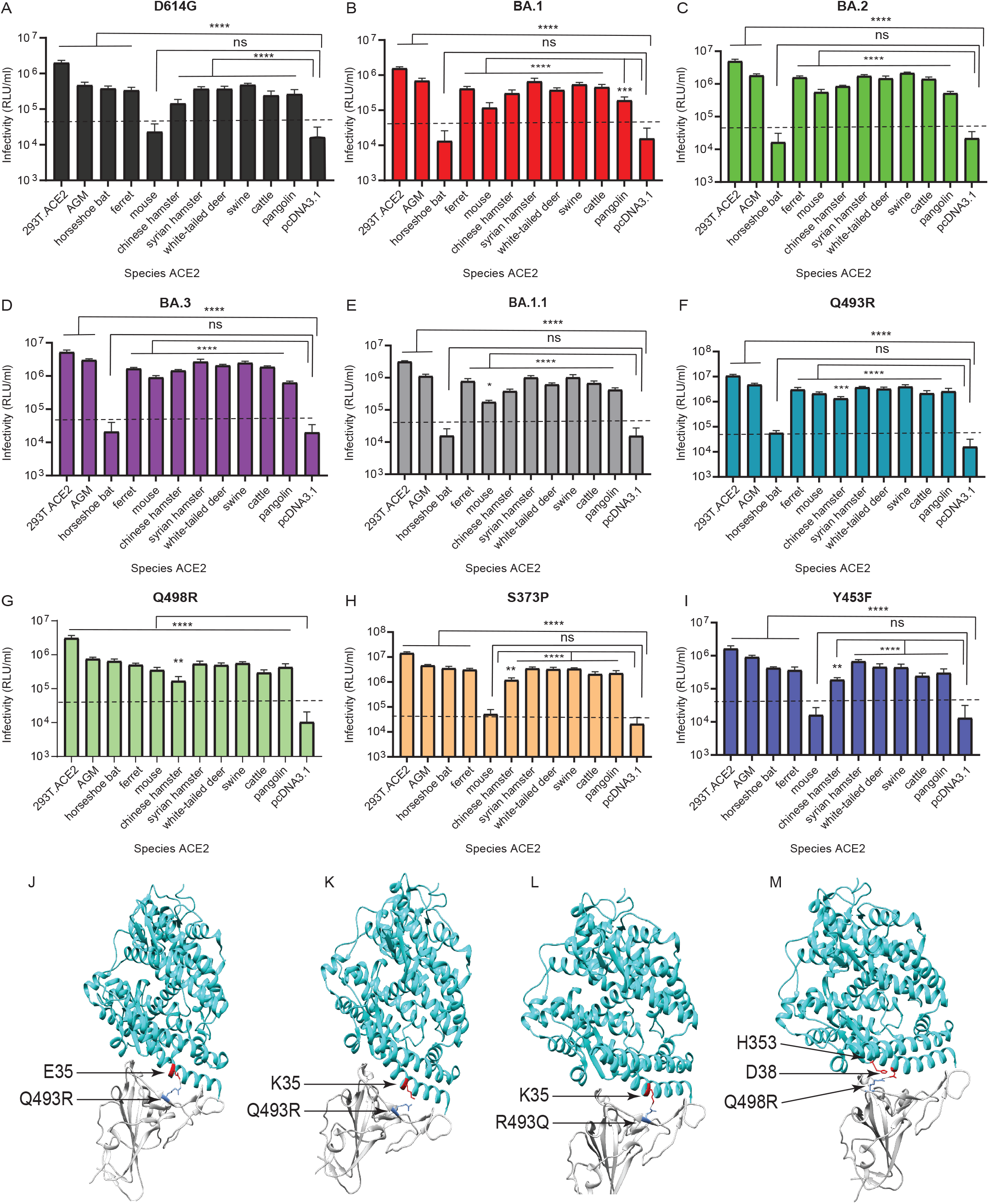
Omicron lineage pseudovirus entry into cells expressing ACE2 from different species. Infectivity of D614G (**A**), BA.1 (**B**) BA.2 (**C**) BA.3 (**D**) BA.1.1 (**E**), Q493R (**F**), Q498R (**G**), S373P (**H**), and Y453F (**I**) pseudoviruses on 293T cells transiently transfected with ACE2 orthologs of indicated species. The ACE2 of African green monkey is denoted as ‘AGM’. 293T cells expressing ACE2 of indicated species, as well as control 293T.ACE2 cells stably expressing human ACE2, were simultaneously infected with 10^6^ RLU/ml of indicated pseudoviruses. Luciferase activities were determined 48 hours post infection. Note: ‘ns’ denotes not significant. Significant differences in infectivity between each species ACE2 compared to pcDNA3.1 control for pseudoviruses are denoted by asterisks, *: *p ≤ 0.05;* **:*p ≤ 0.01;* ***:*p ≤ 0.0005;* ****:*p ≤ 0.0001*. Dotted line indicates background level infection. Results shown are representative of three independent experiments with eight intraassay replicates. The SARS-CoV-2 Omicron RBD-ACE2 interface (PDB: 7WBP) is shown with contacting residues as sticks at the RBD-ACE2 interface (J-M). The SARS-CoV-2 Omicron RBD and ACE2 are colored in grey and cyan respectively. Positions in RBD (blue) that contact ACE2 (red) residues are highlighted. Residue positions are indicated by arrows. The SARS-CoV-2 RBD/ACE2 interactions between 493R/E35 (**J**), R493/K35 (**K**), Q493/K35 (**L**), and R498/D38-H353 (**M**) are shown.

**Table 1:**
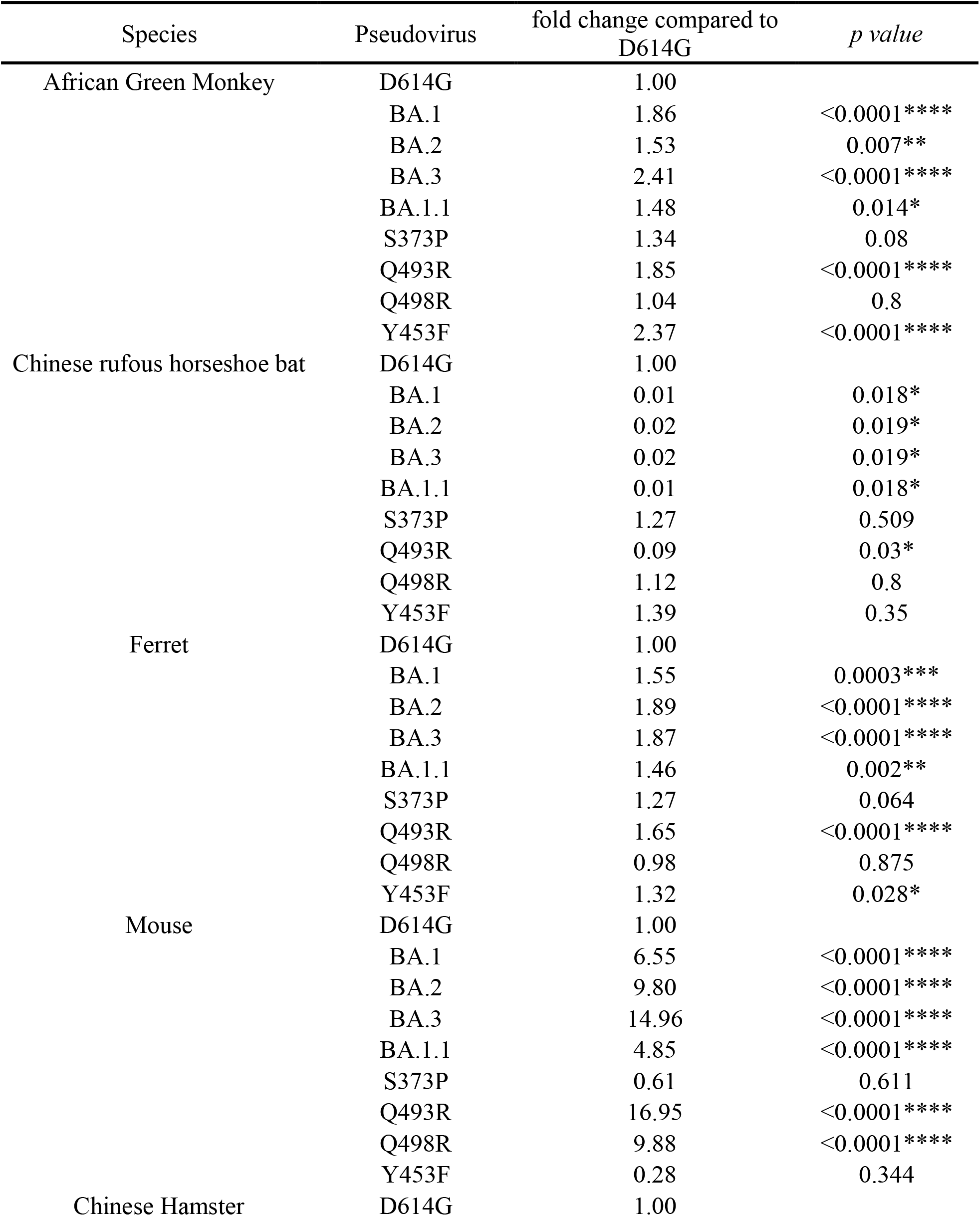
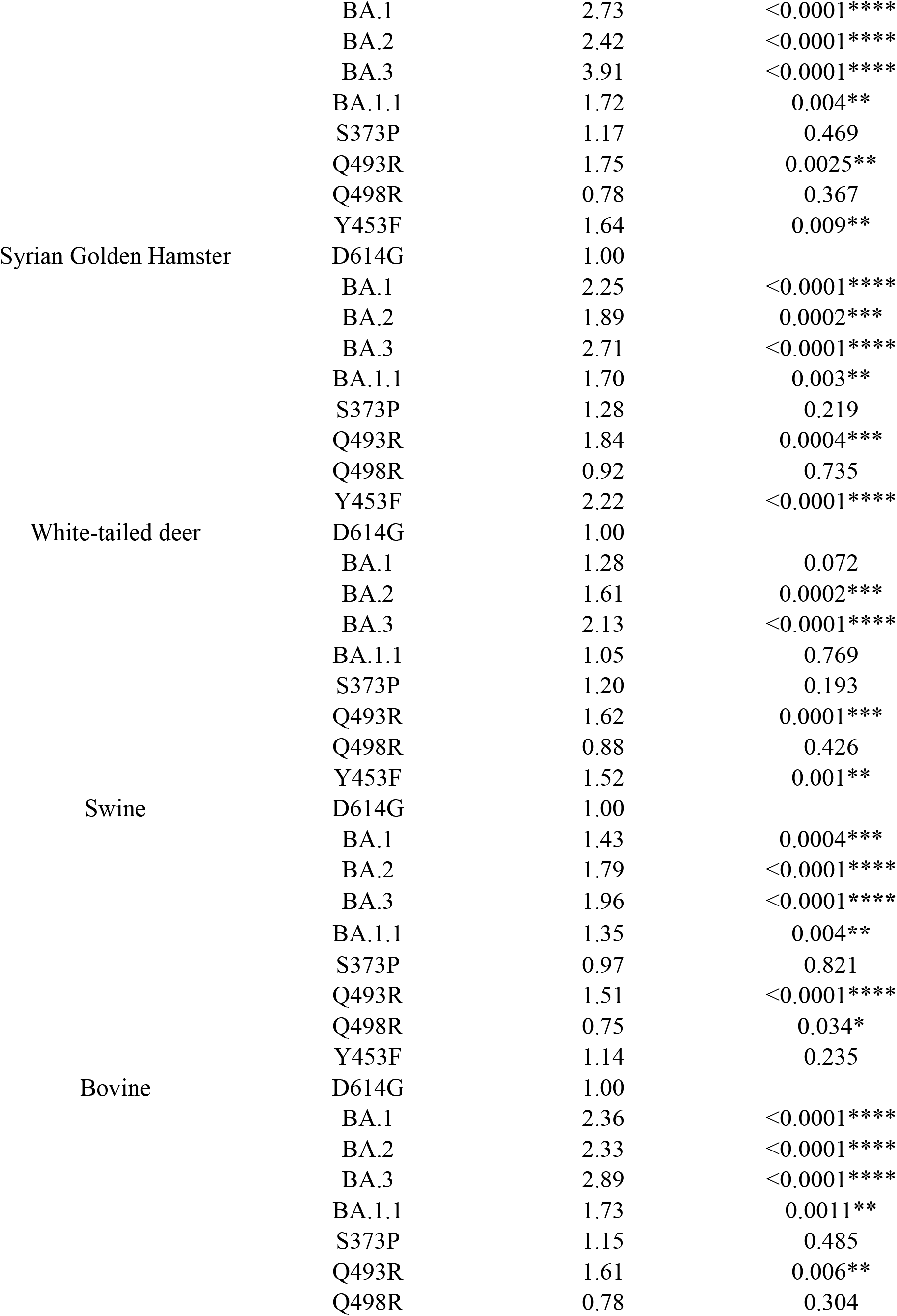
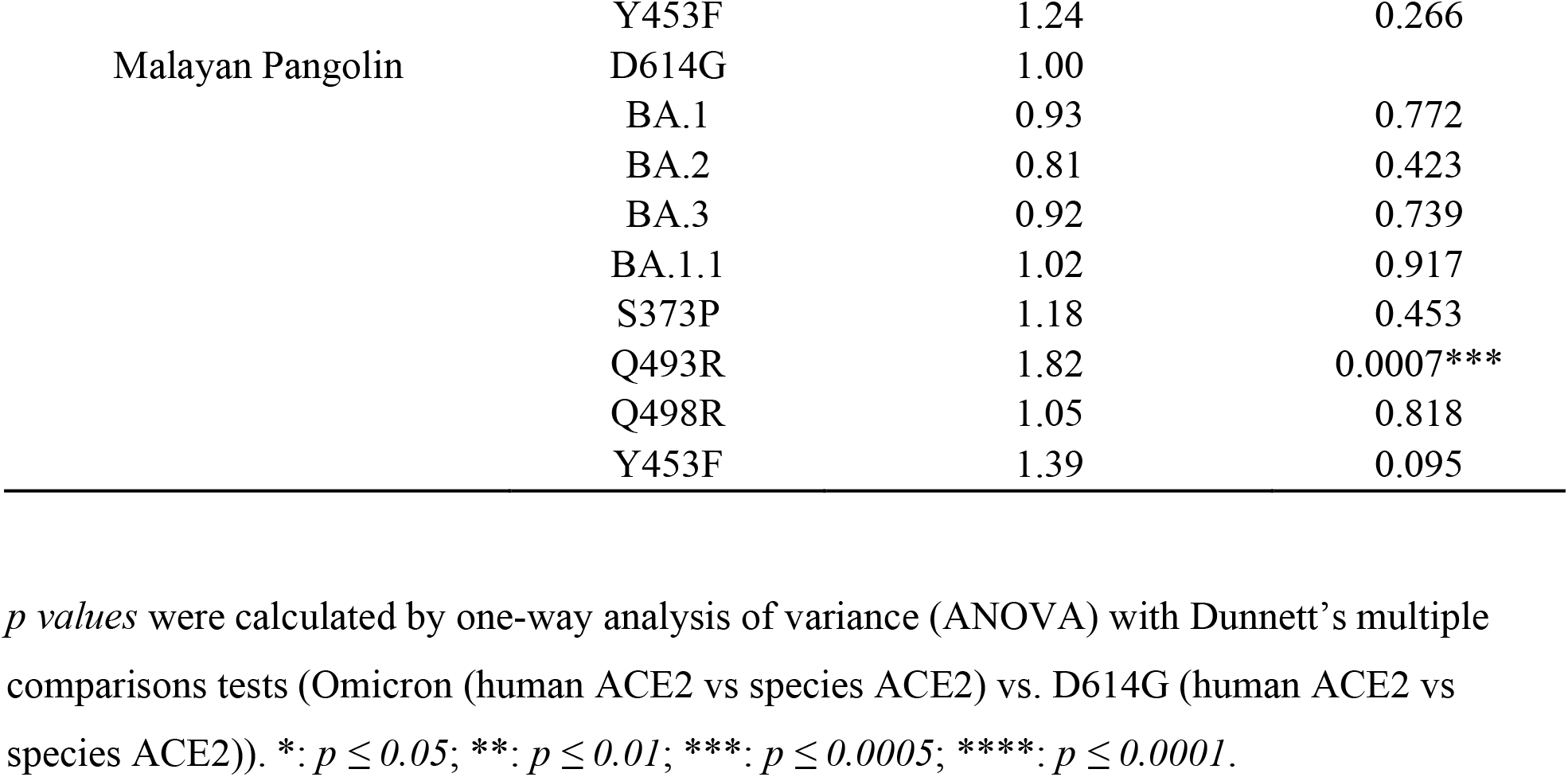
Comparison of relative infectivity (vs infectivity on 293T.ACE2) of pseudoviruses for each species ACE2.

We next investigated the residues in D614G and Omicron spikes that allow or prevent infection of cells expressing horseshoe bat and mouse ACE2, respectively. Crystal structures of SARS-CoV-2 RBD-ACE2 complexes show the ACE2 N-terminal helix cradled in the ridged concave surface formed by the spike RBM ^43–46^. Seventeen RBM residues contact ACE2 residues in its N terminus (Table 2). Among wild type SARS-CoV-2 RBM residues, Q498 interacts with D38, Y41, Q42, L45, and K353 residues of ACE2, while Q493 interacts with K31, H34, and E35 of ACE2, forming a network of hydrogen bonds ^43–46^. Structural and computational determination of Omicron RBD-ACE2 interactions revealed Q493R forming a new salt bridge with E35 while disrupting previous interactions of Q493 with K31 observed in wild type. The Q498R substitution also forms a new hydrogen bond and a salt bridge with ACE2 D38 and Q42 ^47–49^. We assessed whether Q493R and Q498R substitutions in the RBM of Omicron spike facilitate entry of Omicron pseudoviruses in cells expressing mouse ACE2, or conversely, confer resistance to entry of cells expressing horseshoe bat ACE2. Pseudoviruses bearing Q493R and Q498R substitutions on the D614G background spike were therefore generated to infect 293T cells expressing ACE2 proteins of all the species described above. As expected, the Q493R substitution rescued pseudovirus infection in cells expressing mouse ACE2, but reduced pseudovirus infection in cells expressing horseshoe bat ACE2 to background levels (Table 1) (Figure 5F). The Q498R substitution also rescued infection in cells expressing mouse ACE2 above the background but had no effect on cells expressing horseshoe bat ACE2 (Table 1) (Figure 5G). As controls, the Omicron non-RBM S373P and mink adapted Y453F substitutions had no effect on pseudovirus entry in cells expressing mouse ACE2 or horseshoe bat ACE2 (Table 1) (Figure 5H and 5I). It is likely that the Q493R basic substitution in Omicron VOCs forms a salt bridge with the acidic E35 residue of mouse ACE2 or disrupts interaction with the basic K35 residue of horseshoe bat ACE2, thus facilitating or preventing infection of cells expressing mouse ACE2 and horseshoe bat ACE2 respectively (Table 2) (Figure 5J, 5K). Alternatively, the K35 residue in horseshoe bat ACE2 interacts with Q493 residue in D614G spike (Figure 5L). On the other hand, substitutions similar to Q498R were previously observed in mouse-adapted SARS-CoV-2 (Q498H, Q498Y/P499T) to strengthen interaction with D38 of mouse ACE2, which otherwise forms an intramolecular salt bridge with H353 ^50–52^ (Table 2) (Figure 5M). The interactions between R498 in Omicron spike and D38 and H353 in mouse ACE2 likely plays an important role for infection.

**Table 2:** ACE2 residues of species known to interact with SARS-CoV-2 spike receptor binding motif

| ACE2 residue position | 19 | 24 | 27 | 28 | 30 | 31 | 34 | 35 | 37 | 38 | 41 | 42 | 45 | 53 | 79 | 82 | 83 | 90 | 322 | 330 | 353 | 354 | 355 | 357 | 393 |
| --- | --- | --- | --- | --- | --- | --- | --- | --- | --- | --- | --- | --- | --- | --- | --- | --- | --- | --- | --- | --- | --- | --- | --- | --- | --- |
| Human | S | Q | T | F | D | K | H | E | E | D | Y | Q | L | N | L | M | Y | N | N | N | K | G | D | R | R |
| African green monkey | S | Q | T | F | D | K | H | E | E | D | Y | Q | L | N | L | M | Y | N | N | N | K | G | D | R | R |
| Chinese rufous horseshoe bat | S | E | M | F | D | K | T | K | E | D | H | Q | L | N | L | N | Y | N | N | N | K | G | D | R | R |
| Ferret | S | L | T | F | E | K | Y | E | E | E | Y | Q | L | N | H | T | Y | D | N | N | K | R | D | R | R |
| House mouse | S | N | T | F | N | K | Q | E | E | D | Y | Q | L | N | T | S | F | T | H | N | H | G | D | R | R |
| Chinese hamster | S | Q | T | F | D | K | Q | E | E | D | Y | Q | L | N | L | N | Y | N | H | N | K | G | D | R | R |
| Syrian golden hamster | S | Q | T | F | D | K | Q | E | E | D | Y | Q | L | N | L | N | Y | N | Y | N | K | G | D | R | R |
| White-tailed deer | S | Q | T | F | E | K | H | E | E | D | Y | Q | L | N | M | T | Y | N | H | N | K | G | D | R | R |
| Swine | S | L | T | F | E | K | L | E | E | D | Y | Q | L | N | I | T | Y | T | N | N | K | G | D | R | R |
| Bovine | S | Q | T | F | E | K | H | E | E | D | Y | Q | L | N | M | T | Y | N | Y | N | K | G | D | R | R |
| Malayan pangolin | S | E | T | F | E | K | S | E | E | E | Y | Q | L | N | I | N | Y | N | K | N | K | H | D | R | R |

Amino acid substitutions are colored according to its property compared to human residue at similar position; Nonconservative (orange), semiconservative (yellow), or conservative (blue) residues are shown as highlighted.

For the remaining species, S373P, Y453F, Q493R, and Q498R substitutions phenocopied D614G in ACE2 usage (Figure 5F-5I). The Q493R substitution significantly enhanced the usage of all the remainder species, highlighting the role played by this substitution in enhancing ACE2 usage of Omicron variants, as described above (Figure 5F, Table 1). While S373P and Q498R substitutions had no significant effect compared to D614G, the mink-adapted Y453F substitution led to enhanced usage of African green monkey, ferret, Chinese hamster, Syrian golden hamster, white-tailed deer, and swine ACE2 receptors. A previous report has indicated the Y453F substitution enhanced spike interaction with *Mustela* species ACE2 orthologs, including minks, ferrets and stouts, while not compromising human ACE2 usage (Figure 5I, Table 1) ^40^. Our findings extend this report and provide further insights into the effect of Y453F substitution for other species’ ACE2.

### Enhanced neutralization of Omicron variants by vaccinee sera post booster

We next evaluated the serum neutralizing activity in serum from vaccinated individuals who received three immunizations (two dose primary vaccine series and a third dose of booster) of the Pfizer/BNT162b2 vaccine. Receipt of a booster dose elicited significantly higher neutralization titers against D614G (GMT:7527) than two dose primary vaccine series, as seen previously ^53^. Furthermore, all serum samples had neutralizing activity against Omicron pseudoviruses after the receipt of a third booster dose. Neutralization titers against BA.1 (GMT:1087), BA.2 (GMT:961), BA.3 (GMT:916), and BA.1.1 (GMT:916) pseudoviruses were significantly reduced, 6.6-fold (*p≤0.05*), 7.8-fold (*p≤0.0005*), 8.2-fold (*p≤0.0001*), and 7.7-fold (*p≤0.0005*) respectively, compared to D614G (GMT:7527) (Figure 6A).

**Figure 6.**
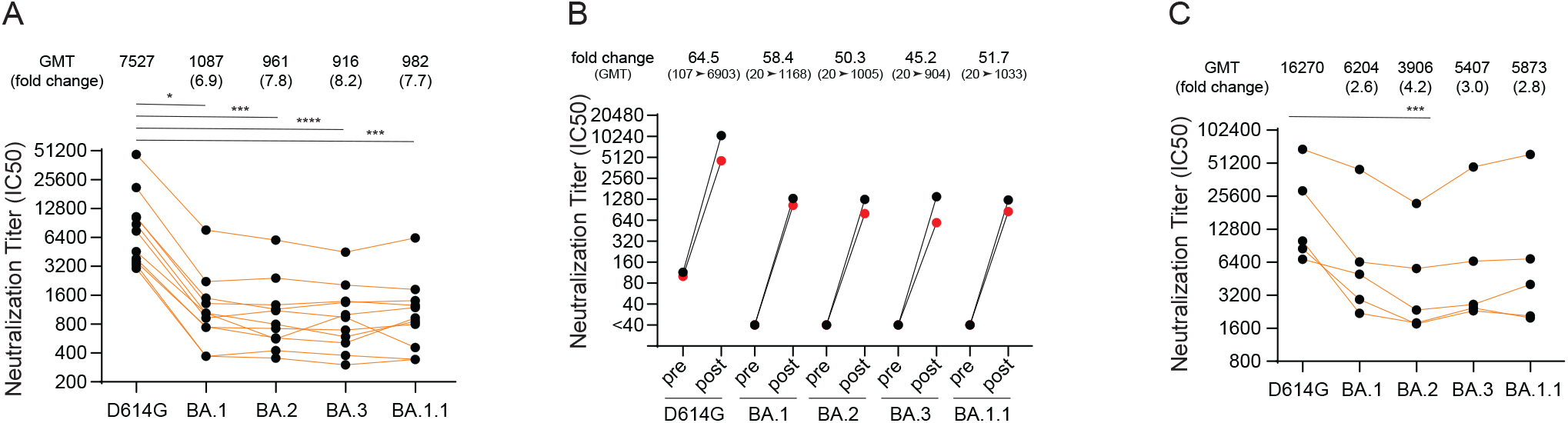
The sensitivity of Omicron (BA) lineage pseudoviruses to post vaccine booster and vaccine-break through infection sera. **A.** Neutralization of D614G and Omicron (BA) lineage pseudoviruses to vaccine-booster-elicited sera obtained from individuals that received primary two-dose series and a third booster dose of Pfizer/BNT162b2 mRNA vaccine. **B.** Sera neutralization titers pre- and post-booster receipt in individuals. **C.** Titers of five vaccine break through cases that experienced Omicron infection. *:*p ≤ 0.05;* **:*p ≤ 0.01;* ***:*p ≤ 0.0005;* ****: *p ≤ 0.0001* compared to D614G. Results shown are from two independent experiments.

For two of the vaccinated individuals described above, we obtained serum samples one week prior to a third booster dose, as well as three weeks after the third booster. Therefore, we compared neutralization activity before and after the booster dose (pre- and post-boost respectively). We found 45.2-64.5-fold enhancement in post-boost neutralization titer compared to pre-boost titer for D614G and Omicron VOCs (Figure 6B).

We also evaluated convalescent sera from five vaccinated individuals who experienced breakthrough infection post boost, when BA.1 or BA.1.1 variants were predominant (December 2021-January 2022). Serum samples from all individuals had the highest neutralization activity against D614G pseudoviruses (GMT: 16270) followed by lower neutralization against BA.1 (GMT: 6204), BA.2 (GMT: 3906), BA.3 (GMT: 5407) and BA.1.1 (GMT: 5873) variant pseudoviruses (Figure 6C). Although the sample size is small, the results show similar resistance trends among these Omicron variants compared to D614G.

## Discussion

In the present study we compared four Omicron lineage (BA.1, BA.2, BA.3 and BA.1.1) variants spikes for their ability to mediate virus entry in cells expressing human ACE2 in the presence or absence of TMPRSS2 and in cells expressing ACE2 orthologs of different species. We also assessed neutralization sensitivity of these spikes by soluble ACE2 and vaccine-elicited antibodies. We found that TMPRSS2 enhances Omicron (BA.1, BA.2, BA.3 and BA.1.1) pseudovirus infection relatively less than the B.1 (D614G) pseudovirus. In addition, compared to D614G, all Omicron lineage spikes more efficiently used ACE2 receptors from diverse animal species including mouse ACE2 but not horseshoe bat ACE2 for entry. Finally, we also observed that three doses of the Pfizer/BNT162b2 vaccine induced antibodies that had comparable levels of neutralization towards BA.1, BA.2, BA.3 and BA.1.1 Omicron lineages, although the neutralization titers against Omicron are 7-8-fold lower than those against D614G.

Several recent studies demonstrated less enhancement of virus entry by TMPRSS2 for BA.1 pseudovirus infection compared to D614G and Delta pseudoviruses in ACE2/TMPRSS2-expressing cell lines (Calu3, Caco2, VeroE6/TMPRSS2), while no infection differences were observed in cell lines expressing ACE2 but not TMPRSS2 (VeroE6, H1299, HeLa-, HEK293T- and A549-ACE2) ^6,54,55^. Similar observations were reported in studies that performed authentic SARS-CoV-2 infections ^6,54,55^. Compared to D614G or the Delta VOC, Omicron variants (BA.1, BA.1.1) had attenuated or inefficient replication in TMPRSS2-positive lower airway or gall bladder organoids, Calu3, and Caco2 lung cell lines, but had no significant difference in cells with low or no TMPRSS2 expression (H1299, HeLa- and 293T-ACE2 cells) ^6,36,55^. Due to apparently less efficient use of TMPRSS2, Omicron variants appear to favor endocytic route of entry rather than TMPRSS2-mediated entry at the cell surface and consequently are more sensitive to endosomal inhibitors (chloroquine, bafilomycin A1 and E64d) ^6,35^. Our findings are consistent with the increased use of the endosomal entry pathway in cells expressing only ACE2, though we found that coexpression of TMPRSS2 enhanced infection.

A possible mechanism that contributes to less efficient TMPRSS2 utilization by Omicron VOCs is the accumulation of multiple additional basic amino acid substitutions, leading to an overall positively charged Omicron spike proteins compared to the D614G spike. These substitutions may confer greater sensitivity to low pH-induced conformational changes in endosomes, and therefore be better adapted for entry in the low pH environment encountered in the upper airway ^6^. In this regard, a recent preprint showed greater sensitivity of Omicron (BA.1, BA.2 and BA.4) pseudoviruses to endosomal (E64d) entry inhibition compared to pseudoviruses with D614G and Delta VOC spikes, but refractory to Camostat inhibition in Caco-2 cells where both endosomal and TMPRSS2-mediated cell surface entry routes are active ^56^. This phenotype is accounted for by the N969K S2 substitution in Omicron VOCs ^56^. However, maximum likelihood fitting models comparing the entry routes of Omicron suggest that Omicron is also efficient in utilizing TMPRSS2 for entry into human nasal epithelial cells ^56^. Alternatively, the BA.1 spike protein was shown to be less efficiently cleaved at the S1/S2 site compared to wild type and Delta VOC in authentic SARS-CoV-2-infected cells, as well as on virions ^6,7,36,55^. Efficient cleavage at the S1/S2 site is known to be required for exposure of the S2’ site for TMPRSS2 processing after ACE2 binding. Substitutions proximal to the furin cleavage site (H655Y, N679K and P681H), as well as substitutions in the S2 subunit, may contribute to less efficient TMPRSS2 cleavage of Omicron spike ^36^. Finally, higher affinities of BA.1 (6-fold) and BA.2 (11-fold) RBD for ACE2 coupled with transitions into the so called two- or three-RBD-up conformations may also contribute to the efficient use of ACE2 ^48,57^. In the present study, we report higher but similar sensitivities for BA.1, BA.2, and BA.1.1 to inhibition by the soluble ACE2 monomer (3-fold vs D614G) while BA.3 and D614G are comparably less sensitive.

Our data further show that Omicron lineage spike proteins more efficiently use ACE2 receptors from diverse animal species for entry. Our findings showing Omicron lineage spikes’ usage of mouse ACE2 for entry extend previous reports showing enhanced binding of BA.1 and BA.2 RBD to mouse ACE2 ^3,57^. Similar (K417N, N501Y, Q493H/R) ^33,50,58^ or closely related (Q498H) ^50,59^ mouse-adapted substitutions previously found in experimentally infected mice are observed in Omicron RBM that may contribute to binding to mouse ACE2. We show that Q493R and Q498R substitutions alone on D614G background confer the ability to use mouse ACE2. On the other hand, the Omicron spike is unable to use Chinese rufous horseshoe bat (*Rhinolophus sinicus*) ACE2 for entry due to the Q493R substitution. These changes in species tropism are likely due to specific Omicron RBM-ACE2 interfacial residues that promote (R493-E35) or disrupt (R493-K35) interactions in mouse or horseshoe bat ACE2, respectively. In addition, the Q493R substitution also significantly enhanced the usage of all other species of ACE2. Altogether, the spike substitutions in the BA.1, BA.2, BA.3 and BA.1.1 Omicron variants permit robust usage of diverse ACE2 orthologues for entry and thus have the potential to broaden the risk of the Omicron variants to infect animal species and spill back to humans.

Despite several amino acids differences in the spikes of the BA.1, BA.2, BA.3 and BA.1.1 variants, three immunizations with the Pfizer/BNT162b2 vaccine elicited high and comparable levels of neutralizing antibodies against BA.1, BA.2, BA.3 and BA.1.1 pseudoviruses, at least for the short term. Since three-dose mRNA vaccine-induced antibodies elicited robust neutralizing antibodies and was shown to protect against severe disease against the BA.1 VOC ^1^, this protection likely extends to emerging BA.2 VOCs. These findings are consistent with several recent studies highlighting enhanced breadth and potency of three dose mRNA vaccine-induced antibody response against Omicron VOCs ^1,3,9–14^. Cross neutralizing antibodies against Omicron were observed post vaccine boost, but not post 2^nd^ vaccination ^1^, suggesting recalled memory B cell or *de novo* induction of novel B cell clones that permit cross neutralization ^15,16^.

Breakthrough infections with the BA.1 VOC after three vaccinations induced high neutralization titers against all Omicron variant pseudoviruses, as was the case for after three doses of the Pfizer/BNT162b2 vaccine without breakthrough infection. Comparable marked enhancement of serum-neutralizing activity between three dose-vaccinated subjects, vaccine breakthrough, and infected/vaccinated cases was reported in an earlier study where the number of exposures and/or time period between exposures to SARS-CoV-2, either via vaccination or infection, correlated with the strength of neutralizing antibody responses, as well as resilience to variants ^10^. Furthermore, our findings extend a recent report where sera from vaccinated individuals with confirmed Omicron breakthrough infection showed higher neutralization titers compared to vaccinated individuals without breakthrough infection ^60^. Altogether, findings from us and the others suggest that breakthrough infections can boost preexisting immunity induced by three doses of the Pfizer/BNT162b2 vaccine, thereby eliciting antibodies that neutralize not only Omicron and B.1 variants, but also Alpha, Beta, and Delta variants ^60^. Finally, a recent study used homologous hamster sera and antigenic cartography to visualize antigenic evolution of SARS-CoV-2 VOCs, demonstrating distinct antigenicity of BA.1 and BA.2 VOCs, separate from ancestral and earlier SARS-CoV-2 VOCs ^61^.

Our study has several caveats, including the use of pseudoviruses instead of authentic SARS-CoV-2 for conducting experiments. However, our findings using pseudoviruses agree with those reported using authentic SARS-CoV-2. For instance, authentic BA.1/BA.1.1 VOCs were shown to undergo attenuated replication in TMPRSS2-expressing cells compared to ancestral Wuhan-Hu-1, and Alpha, Beta, and Delta VOCs ^6,36^. These reports also showed greater sensitivity of BA.1 pseudovirus entry to endosomal inhibitor E64d. While we used pseudovirus entry assays to determine Omicron variant usage of ACE2 receptors of various animal species, it remains unknown whether there may be intrinsic and/or innate host-specific factors that might act to inhibit live Omicron VOCs at an entry or post entry step. Furthermore, although we identified RBM substitutions in Omicron spike that conferred the ability to use mouse or horseshoe bat ACE2, we didn’t confirm ACE2 substitutions that permit or prevent Omicron spike binding. For instance, introducing K35E substitution in horseshoe bat ACE2 should permit Omicron variants’ usage. Finally, analysis of a limited number of serum samples and short follow up after the receipt of three doses of the Pfizer/BNT162b2 mRNA vaccine do not give us insights into the durability of the antibody response. While studies of antibody durability are ongoing, our findings indicate that three dose immunization with the Pfizer/BNT162b2 will likely contribute to protection from severe disease caused by the ongoing BA.2 VOC.

## Materials and Methods

### Ethics Statement

Sera were obtained from participants who received three doses of the Pfizer/BNT162b2 vaccine and had no serological evidence of SARS-CoV-2 infection prior to vaccination. The first two doses of the Pfizer/BNT162b2 vaccine were received before March 1, 2021, whereas the third dose of Pfizer/BNTech162b2 vaccine was received by December 15, 2021. Sera were also obtained from five individuals who experienced vaccine breakthrough infection between December 2021 and January 2022, when BA.1 or BA.1.1 VOCs were dominant. Sera were collected at the U.S. Food and Drug Administration with written consent under an approved Institutional Review Board (IRB) protocol (FDA IRB Study # 2021-CBER-045).

### Plasmids and Cell Lines

Codon-optimized, full-length open reading frames of the spike genes of B.1 (D614G) and Omicron variants in the study were synthesized into pVRC8400 (B.1, BA.1, BA.2, and BA.3) or pcDNA3.1(+) (BA.1.1) were obtained from the Vaccine Research Center (National Institutes of Health, Bethesda, MD) and GenScript (Piscataway, NJ, USA). The codon optimization parameters for spike gene expression in human cells follow GenScript’s Optimum Gene algorithm as described previously ^53^. The spike substitutions present in the Omicron variants spikes are listed in Figure 1. The HIV gag/pol packaging (pCMVΔR8.2) and firefly luciferase encoding transfer vector (pHR’CMV-Luc) plasmids ^62,63^ were obtained from the Vaccine Research Center (National Institutes of Health, Bethesda, MD, USA). ACE2 genes of various species (African green monkey (AGM), Chinese rufous horseshoe bat (*Rhinolophus sinicus*), ferret, mouse, Chinese hamster, Syrian golden hamster, white-tailed deer, swine, bovine, and pangolin) with a C-terminal V5 tag were synthesized by GenScript as described previously ^42^. 293T (ATCC, Manassas, VA, USA; Cat no: CRL-11268), 293T.ACE2 (BEI Resources, Manassas, VA, USA; Cat no: NR-52511) ^64^ and 293T.ACE2.TMPRSS2 cells stably expressing human angiotensin-converting enzyme 2 (ACE2) and transmembrane serine protease 2 (TMPRSS2) (BEI Resources, Manassas, VA, USA; Cat no: NR-55293) ^34^ were maintained at 37°C in Dulbecco’s modified eagle medium (DMEM) supplemented with high glucose, L-glutamine, minimal essential media (MEM) non-essential amino acids, penicillin/streptomycin, HEPES, and 10% fetal bovine serum (FBS).

### SARS-CoV-2 Pseudovirus Production and Neutralization Assay

HIV-based lentiviral pseudoviruses with desired spike proteins (D614G, BA.1, BA.2, BA.3, and BA.1.1) were generated as previously described ^34,65^. Pseudoviruses comprising the spike glycoprotein and a firefly luciferase (FLuc) reporter gene packaged within HIV capsid were produced in 293T cells by co-transfection of 5 μg of pCMVΔR8.2, 5 μg of pHR’CMVLuc and 0.5 μg of pVRC8400 or 4 μg of pcDNA3.1(+) encoding a codon-optimized spike gene. Pseudovirus supernatants were collected approximately 48 h post transfection, filtered through a 0.45 μm low protein binding filter, and stored at −80°C.

Neutralization assays were performed using 293T.ACE2.TMPRSS2 cells in 96-well plates as previously described ^34,65^. Pseudoviruses with titers of approximately 10^6^ relative luminescence units per milliliter (RLU/mL) of luciferase activity were incubated with serially diluted sera or inhibitors for two hours at 37°C prior to inoculation onto the plates that were preseeded one day earlier with 3.0 × 10^4^ cells/well. Pseudovirus infectivity was determined 48 h post inoculation for luciferase activity by luciferase assay reagent (Promega) according to the manufacturer’s instructions. The inhibitor concentration or inverse of the sera dilutions causing a 50% reduction of RLU compared to control was reported as the neutralization titer. Titers were calculated using a nonlinear regression curve fit (GraphPad Prism Software Inc., La Jolla, CA, USA). The mean titer from at least two independent experiments each with intra-assay duplicates was reported as the final titer. For experiments involving camostat mesylate (0.03–500 μM) and chloroquine (0.39-25 μM) inhibitors, each target cell type was pretreated with inhibitor for two hours before pseudovirus infection in the presence of respective inhibitor as described previously ^34^.

### Soluble ACE2 Protein Production

His-tagged soluble human ACE2 was produced in FreeStyle™ 293-F cells by transfecting soluble human ACE2 (1-741 aa) expression vector plasmid DNA using 293fectin (Thermo Fisher) and purified using HiTrap Chelating column charged with nickel (GE healthcare) according to the manufacturer’s instructions. The eluate containing soluble ACE2 was concentrated to 1.0 mL. Protein containing fractions were pooled and concentrated using Amicon ultra-15 ultracentrifugal unit. The purified proteins were analyzed on a 4%–12% SDS-PAGE stained with Coomassie blue, or membrane probed with mouse monoclonal 6x-His tag antibody (4A12E4) (Thermofisher, Waltham, MA) (Supplementary Figure 2).

### Soluble ACE2 Neutralization Using SARS-CoV-2 Pseudoviruses

Soluble human ACE2 neutralization assays were performed using 293T.ACE2.TMPRSS2 as previously described ^53^. Briefly, pseudoviruses were treated with 3-fold serial dilutions of soluble ACE2 for one hour at 37°C. Pseudovirus and soluble ACE2 mixtures (100 μl) were then inoculated onto 96-well plates that were pre-seeded with 3.0 x 10^4^ cells per well one day prior to the assay. Pseudovirus firefly luciferase activity was determined 48 h post inoculation. The ACE2 concentration causing a 50% reduction of luciferase activity compared to untreated control was reported as the IC_50_ using a nonlinear regression curve fit (GraphPad Prism software Inc., La Jolla, CA).

### Western Blotting

Cell lysates were resuspended in 1X Laemmli loading buffer containing 2-mercaptoethanol, heated at 70°C for 10 min, resolved by 4–20% SDS PAGE and transferred onto nitrocellulose membranes. Membranes were probed for the V5-tag and γ-actin using V5 epitope tag antibody (Novus Biologicals, Centennial, CO), and mouse gamma actin polyclonal antibody (Thermofisher), respectively.

### Computational Analysis

Contact residues in Omicron RBD/human ACE2 complexes are shown as sticks on the Protein Data Bank entry (PDB) code 7WBP ^47^ by UCSF Chimera program (http://www.cgl.ucsf.edu/chimera/). Substitutions in RBD/ACE2 complex were introduced by rotamers function of USCF Chimera program.

### Statistical Analyses

One-way analysis of variance (ANOVA) with Dunnett’s multiple comparisons tests (Omicron variants compared to D614G), and geometric mean titers (GMT) with 95% confidence intervals were performed using GraphPad Prism software. The *p* values of less than 0.05 were considered statistically significant. All neutralization titers were log_2_ transformed for analyses.

---

**Supplementary figure 1.**
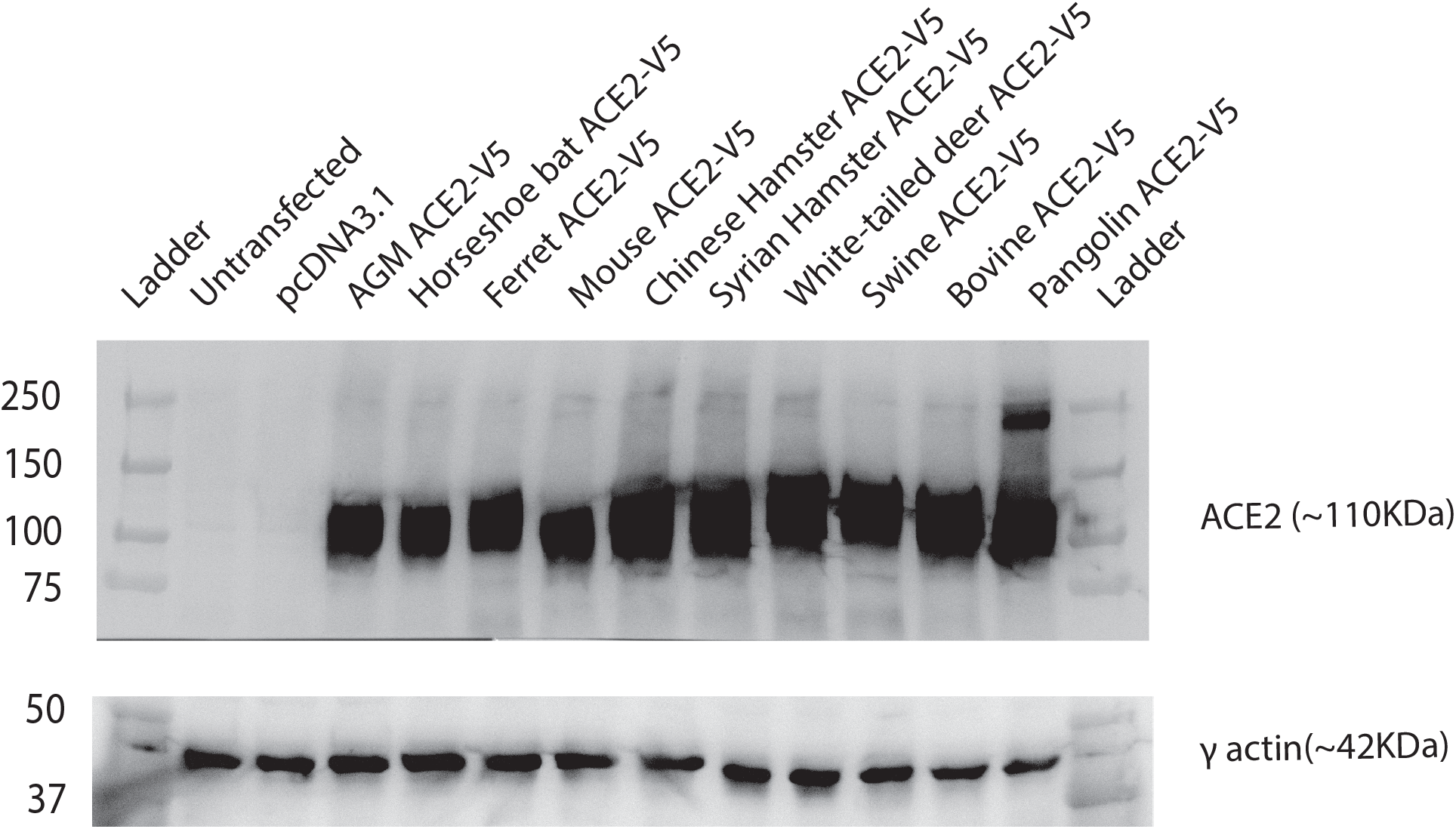
Transit expression of variant species ACE2. Western blotting of V5-tagged ACE2 proteins belonging to different species using Anti-V5 Tag antibody. Cellular control is γ-actin.

**Supplementary figure 2.**
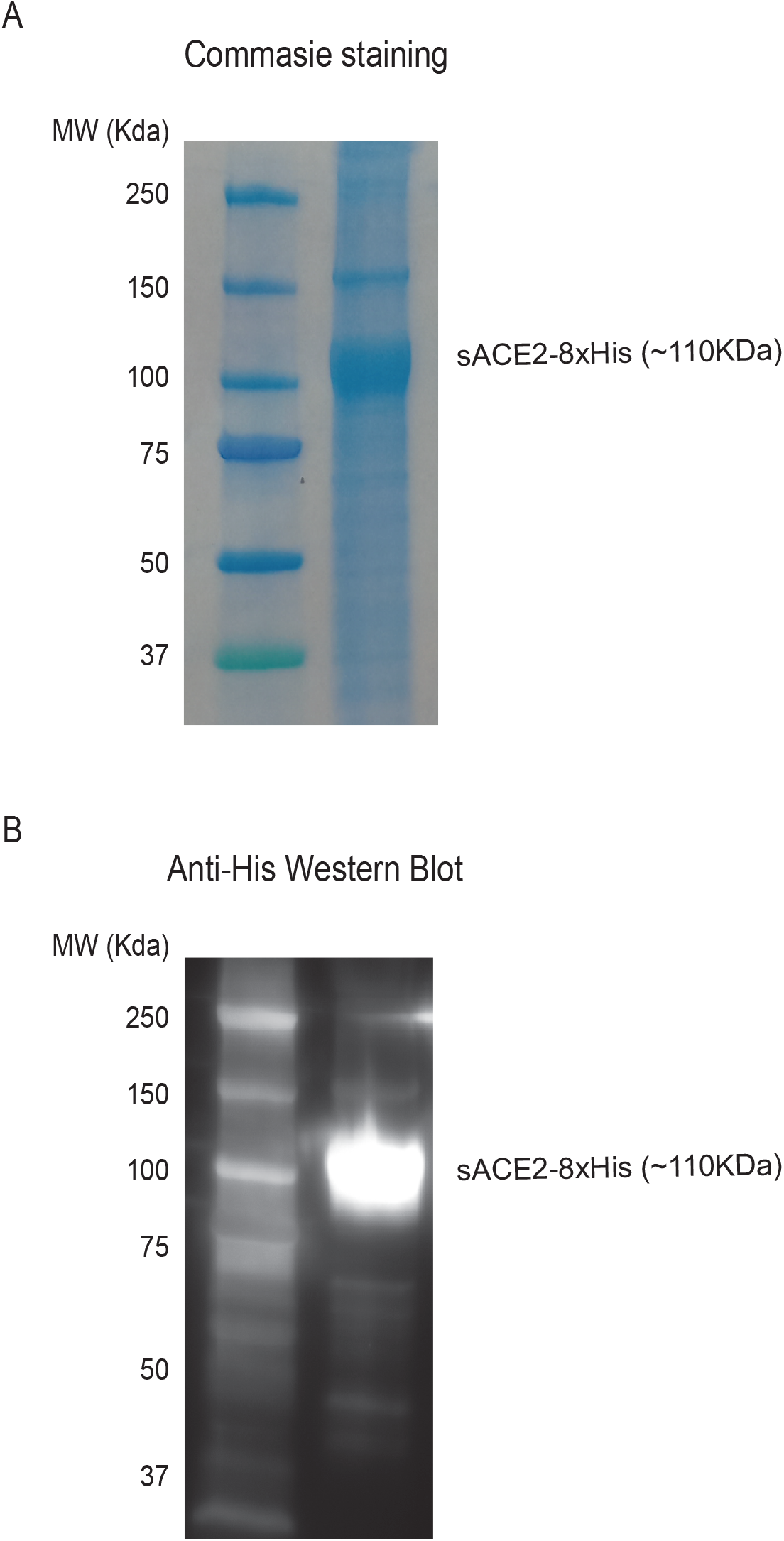
The detection of purified soluble human ACE2. **A.** Commassie staining of purified soluble ACE2 (sACE2-8XHis) protein. **B.** Western blotting of sACE2-8XHis protein using Anti-6xHis Tag antibody

## References

1. Lusvarghi S, Pollett SD, Neerukonda SN, et al. SARS-CoV-2 BA.1 variant is neutralized by vaccine booster-elicited serum, but evades most convalescent serum and therapeutic antibodies. Science Translational Medicine. 0(0):eabn8543.

2. Cele S, Jackson L, Khoury DS, et al. Omicron extensively but incompletely escapes Pfizer BNT162b2 neutralization. Nature. 2022;602(7898):654–656.

3. Cameroni E, Bowen JE, Rosen LE, et al. Broadly neutralizing antibodies overcome SARS-CoV-2 Omicron antigenic shift. Nature. 2022;602(7898):664–670.

4. Schmidt F, Muecksch F, Weisblum Y, et al. Plasma Neutralization of the SARS-CoV-2 Omicron Variant. New England Journal of Medicine. 2021;386(6):599–601.

5. Wilhelm A, Widera M, Grikscheit K, et al. Reduced Neutralization of SARS-CoV-2 Omicron Variant by Vaccine Sera and monoclonal antibodies. medRxiv. 2021:2021.2012.2007.21267432.

6. Meng B, Abdullahi A, Ferreira IATM, et al. Altered TMPRSS2 usage by SARS-CoV-2 Omicron impacts infectivity and fusogenicity. Nature. 2022;603(7902):706–714.

7. Yamasoba D, Kimura I, Nasser H, et al. Virological characteristics of SARS-CoV-2 BA.2 variant. bioRxiv. 2022:2022.2002.2014.480335.

8. Suzuki R, Yamasoba D, Kimura I, et al. Attenuated fusogenicity and pathogenicity of SARS-CoV-2 Omicron variant. Nature. 2022;603(7902):700–705.

9. Sheward DJ, Kim C, Ehling RA, et al. Neutralisation sensitivity of the SARS-CoV-2 omicron (B.1.1.529) variant: a cross-sectional study. The Lancet Infectious Diseases. 2022.

10. Walls AC, Sprouse KR, Bowen JE, et al. SARS-CoV-2 breakthrough infections elicit potent, broad, and durable neutralizing antibody responses. Cell. 2022;185(5):872–880.e873.

11. Planas D, Saunders N, Maes P, et al. Considerable escape of SARS-CoV-2 Omicron to antibody neutralization. Nature. 2022;602(7898):671–675.

12. Garcia-Beltran WF, St. Denis KJ, Hoelzemer A, et al. mRNA-based COVID-19 vaccine boosters induce neutralizing immunity against SARS-CoV-2 Omicron variant. Cell. 2022;185(3):457–466.e454.

13. Pérez-Then E, Lucas C, Monteiro VS, et al. Neutralizing antibodies against the SARS-CoV-2 Delta and Omicron variants following heterologous CoronaVac plus BNT162b2 booster vaccination. Nature Medicine. 2022;28(3):481–485.

14. Bowen JE, Sprouse KR, Walls AC, et al. Omicron BA.1 and BA.2 neutralizing activity elicited by a comprehensive panel of human vaccines. bioRxiv. 2022:2022.2003.2015.484542.

15. Goel RR, Painter MM, Lundgreen KA, et al. Efficient recall of Omicron-reactive B cell memory after a third dose of SARS-CoV-2 mRNA vaccine. bioRxiv. 2022:2022.2002.2020.481163.

16. Muecksch F, Wang Z, Cho A, et al. Increased Potency and Breadth of SARS-CoV-2 Neutralizing Antibodies After a Third mRNA Vaccine Dose. bioRxiv. 2022:2022.2002.2014.480394.

17. Xia H, Zou J, Kurhade C, et al. Neutralization and durability of 2 or 3 doses of the BNT162b2 vaccine against Omicron SARS-CoV-2. Cell Host & Microbe. 2022.

18. Vanshylla K, Tober-Lau P, Gruell H, et al. Durability of omicron-neutralising serum activity after mRNA booster immunisation in older adults. The Lancet Infectious Diseases. 2022;22(4):445–446.

19. Oreshkova N, Molenaar RJ, Vreman S, et al. SARS-CoV-2 infection in farmed minks, the Netherlands, April and May 2020. Eurosurveillance. 2020;25(23):2001005.

20. Ritter JM, Wilson TM, Gary JM, et al. Histopathology and localization of SARS-CoV-2 and its host cell entry receptor ACE2 in tissues from naturally infected US-farmed mink (Neovison vison). Veterinary Pathology.0(0):03009858221079665.

21. Zhang Q, Zhang H, Gao J, et al. A serological survey of SARS-CoV-2 in cat in Wuhan. Emerging Microbes & Infections. 2020;9(1):2013–2019.

22. Patterson EI, Elia G, Grassi A, et al. Evidence of exposure to SARS-CoV-2 in cats and dogs from households in Italy. Nature Communications. 2020;11(1):6231.

23. Račnik J, Kočevar A, Slavec B, et al. Transmission of SARS-CoV-2 from Human to Domestic Ferret. Emerg Infect Dis. 2021;27(9):2450–2453.

24. Koeppel KN, Mendes A, Strydom A, Rotherham L, Mulumba M, Venter M. SARS-CoV-2 Reverse Zoonoses to Pumas and Lions, South Africa. Viruses. 2022;14(1):120.

25. Chandler Jeffrey C, Bevins Sarah N, Ellis Jeremy W, et al. SARS-CoV-2 exposure in wild white-tailed deer (Odocoileus virginianus). Proceedings of the National Academy of Sciences. 2021;118(47):e2114828118.

26. Ulrich L, Wernike K, Hoffmann D, Mettenleiter TC, Beer M. Experimental Infection of Cattle with SARS-CoV-2. Emerg Infect Dis. 2020;26(12):2979–2981.

27. Bosco-Lauth AM, Walker A, Guilbert L, et al. Susceptibility of livestock to SARS-CoV-2 infection. Emerging Microbes & Infections. 2021;10(1):2199–2201.

28. Pickering BS, Smith G, Pinette MM, et al. Susceptibility of Domestic Swine to Experimental Infection with Severe Acute Respiratory Syndrome Coronavirus 2. Emerg Infect Dis. 2021;27(1):104–112.

29. Gaudreault NN, Cool K, Trujillo JD, et al. Susceptibility of sheep to experimental coinfection with the ancestral lineage of SARS-CoV-2 and its alpha variant. Emerging Microbes & Infections. 2022;11(1):662–675.

30. Bao L, Deng W, Huang B, et al. The pathogenicity of SARS-CoV-2 in hACE2 transgenic mice. Nature. 2020;583(7818):830–833.

31. Jiang R-D, Liu M-Q, Chen Y, et al. Pathogenesis of SARS-CoV-2 in Transgenic Mice Expressing Human Angiotensin-Converting Enzyme 2. Cell. 2020;182(1):50–58.e58.

32. Sun S-H, Chen Q, Gu H-J, et al. A Mouse Model of SARS-CoV-2 Infection and Pathogenesis. Cell Host & Microbe. 2020;28(1):124–133.e124.

33. Leist SR, Dinnon KH, Schäfer A, et al. A Mouse-Adapted SARS-CoV-2 Induces Acute Lung Injury and Mortality in Standard Laboratory Mice. Cell. 2020;183(4):1070–1085.e1012.

34. Neerukonda SN, Vassell R, Herrup R, et al. Establishment of a well-characterized SARS-CoV-2 lentiviral pseudovirus neutralization assay using 293T cells with stable expression of ACE2 and TMPRSS2. PLOS ONE. 2021;16(3):e0248348.

35. Zhao H, Lu L, Peng Z, et al. SARS-CoV-2 Omicron variant shows less efficient replication and fusion activity when compared with Delta variant in TMPRSS2-expressed cells. Emerging Microbes & Infections. 2022;11(1):277–283.

36. Shuai H, Chan JF-W, Hu B, et al. Attenuated replication and pathogenicity of SARS-CoV-2 B.1.1.529 Omicron. Nature. 2022;603(7902):693–699.

37. Wang Q, Anang S, Iketani S, et al. Functional properties of the spike glycoprotein of the emerging SARS-CoV-2 variant B.1.1.529. bioRxiv. 2021:2021.2012.2027.474288.

38. Wang Q, Nair MS, Anang S, et al. Functional differences among the spike glycoproteins of multiple emerging severe acute respiratory syndrome coronavirus 2 variants of concern. iScience. 2021;24(11):103393.

39. Zhou J, Peacock TP, Brown JC, et al. Mutations that adapt SARS-CoV-2 to mink or ferret do not increase fitness in the human airway. Cell Reports. 2022;38(6):110344.

40. Ren W, Lan J, Ju X, et al. Mutation Y453F in the spike protein of SARS-CoV-2 enhances interaction with the mink ACE2 receptor for host adaption. PLoS Pathog. 2021;17(11):e1010053–e1010053.

41. Marques AD, Sherrill-Mix S, Everett JK, et al. Evolutionary Trajectories of SARS-CoV-2 Alpha and Delta Variants in White-Tailed Deer in Pennsylvania. medRxiv. 2022:2022.2002.2017.22270679.

42. Liu S, Selvaraj P, Lien CZ, et al. The PRRA Insert at the S1/S2 Site Modulates Cellular Tropism of SARS-CoV-2 and ACE2 Usage by the Closely Related Bat RaTG13. Journal of Virology. 2021;95(11):e01751–01720.

43. Wang Q, Zhang Y, Wu L, et al. Structural and Functional Basis of SARS-CoV-2 Entry by Using Human ACE2. Cell. 2020;181(4):894–904.e899.

44. Yan R, Zhang Y, Li Y, Xia L, Guo Y, Zhou Q. Structural basis for the recognition of SARS-CoV-2 by full-length human ACE2. Science. 2020;367(6485):1444–1448.

45. Lan J, Ge J, Yu J, et al. Structure of the SARS-CoV-2 spike receptor-binding domain bound to the ACE2 receptor. Nature. 2020;581(7807):215–220.

46. Shang J, Ye G, Shi K, et al. Structural basis of receptor recognition by SARS-CoV-2. Nature. 2020;581(7807):221–224.

47. Han P, Li L, Liu S, et al. Receptor binding and complex structures of human ACE2 to spike RBD from omicron and delta SARS-CoV-2. Cell. 2022;185(4):630–640.e610.

48. Hoffmann M, Zhang L, Pöhlmann S. Omicron: Master of immune evasion maintains robust ACE2 binding. Signal Transduction and Targeted Therapy. 2022;7(1):118.

49. Yin W, Xu Y, Xu P, et al. Structures of the Omicron spike trimer with ACE2 and an anti-Omicron antibody. Science. 2022;375(6584):1048–1053.

50. Huang K, Zhang Y, Hui X, et al. Q493K and Q498H substitutions in Spike promote adaptation of SARS-CoV-2 in mice. EBioMedicine. 2021;67:103381.

51. Dinnon KH, Leist SR, Schäfer A, et al. A mouse-adapted model of SARS-CoV-2 to test COVID-19 countermeasures. Nature. 2020;586(7830):560–566.

52. Gawish R, Starkl P, Pimenov L, et al. ACE2 is the critical in vivo receptor for SARS-CoV-2 in a novel COVID-19 mouse model with TNF-and IFNγ-driven immunopathology. eLife. 2022;11:e74623.

53. Neerukonda SN, Vassell R, Lusvarghi S, et al. SARS-CoV-2 Delta Variant Displays Moderate Resistance to Neutralizing Antibodies and Spike Protein Properties of Higher Soluble ACE2 Sensitivity, Enhanced Cleavage and Fusogenic Activity. Viruses. 2021;13(12):2485.

54. Hoffmann M, Krüger N, Schulz S, et al. The Omicron variant is highly resistant against antibody-mediated neutralization: Implications for control of the COVID-19 pandemic. Cell. 2022;185(3):447–456.e411.

55. Zeng C, Evans JP, Qu P, et al. Neutralization and Stability of SARS-CoV-2 Omicron Variant. bioRxiv. 2021:2021.2012.2016.472934.

56. Peacock TP, Brown JC, Zhou J, et al. The altered entry pathway and antigenic distance of the SARS-CoV-2 Omicron variant map to separate domains of spike protein. bioRxiv. 2022:2021.2012.2031.474653.

57. Xu Y, Wu C, Cao X, et al. Structural and biochemical mechanism for increased infectivity and immune evasion of Omicron BA.1 and BA.2 variants and their mouse origins. bioRxiv. 2022:2022.2004.2012.488075.

58. Sun S, Gu H, Cao L, et al. Characterization and structural basis of a lethal mouse-adapted SARS-CoV-2. Nature Communications. 2021;12(1):5654.

59. Zhang Y, Huang K, Wang T, et al. SARS-CoV-2 Rapidly Adapts in Aged BALB/c Mice and Induces Typical Pneumonia. Journal of Virology. 2021;95(11):e02477–02420.

60. Suryawanshi RK, Chen IP, Ma T, et al. Limited cross-variant immunity from SARS-CoV-2 Omicron without vaccination. Nature. 2022.

61. Mykytyn AZ, Rissmann M, Kok A, et al. Omicron BA.1 and BA.2 are antigenically distinct SARS-CoV-2 variants. bioRxiv. 2022:2022.2002.2023.481644.

62. Naldini L, Blömer U, Gallay P, et al. In Vivo Gene Delivery and Stable Transduction of Nondividing Cells by a Lentiviral Vector. Science. 1996;272(5259):263–267.

63. Zufferey R, Nagy D, Mandel RJ, Naldini L, Trono D. Multiply attenuated lentiviral vector achieves efficient gene delivery in vivo. Nat Biotechnol. 1997;15(9):871–875.

64. Crawford KHD, Eguia R, Dingens AS, et al. Protocol and Reagents for Pseudotyping Lentiviral Particles with SARS-CoV-2 Spike Protein for Neutralization Assays. Viruses. 2020;12(5):513.

65. Neerukonda SN, Vassell R, Weiss CD, Wang W. Measuring Neutralizing AntibodiesNeutralizing antibodies to SARS-CoV-2 Using Lentiviral Spike-Pseudoviruses. In: Chu JJH, Ahidjo BA, Mok CK, eds. SARS-CoV-2: Methods and Protocols. New York, NY: Springer US; 2022:305–314.

